# Environmental disorder can tip the population genetics of range expansions

**DOI:** 10.1101/494955

**Authors:** Matti Gralka, Oskar Hallatschek

**Affiliations:** Department of Physics, University of California, Berkeley; Department of Integrative Biology, University of California, Berkeley

## Abstract

Evolutionary dynamics is fundamentally shaped by stochastic processes: spontaneous mutations enter populations randomly, and the fate of a mutant lineage is determined by the competition between (random) genetic drift and (deterministic) selection. In populations undergoing range expansions, fluctuations in the reproductive process and the local motion of individuals are enhanced within a small subpopulation at the edge of the population. Range expansions are typically studied in homogeneous environments, but we argue here that the fluctuations at the range edge are susceptible to small-scale environmental heterogeneities that may have a strong effect on the evolutionary dynamics of the expanding population.

To show this, we tracked the dynamics of the clones of spontaneous mutations with a tunable fitness effect in bacterial colonies grown on randomly disordered surfaces. We find that environmental heterogeneity on scales much larger than an individual, but much smaller than the total population, can dramatically reduce the efficacy of selection. Time lapse microscopy and computer simulations suggest that this effect is a general consequence of a local “pinning” of the expansion front, whereby stretches of the front are slowed down on a length scale that depends on the structure of the environmental heterogeneity. This pinning focuses the range expansion into a small number of individuals with access to expansion paths, increasing the importance of chance and thus limiting the efficacy of selection.

## Introduction

Noise, and its competition with deterministic forces, plays an integral role in biology, such as in stochastic gene expression, cellular decision making, and cell differentiation [2]. Stochasticity is also a crucial component of evolutionary dynamics: not only do the mutations entering a population occur at random times in random individuals and at random positions in their genome, but in addition the fate of a mutation and its clonal lineage is largely stochastic and only partly determined by its effect on the individual’s fitness.

The random fluctuations in the frequency of a mutant allele due to the stochasticity associated with reproduction are called genetic drift. Genetic drift is particularly strong at the front of range expansions because only a relatively small number of individuals at the front of the expansion contributes to future growth and thus has any influence on the future genotypic composition of the population. The neutral diversity and adaptation in spatially expanding populations has been studied in computer simulations [11,33], in the field [38,44,50], and in microbial colonies [16,21,24,35], where nutrient gradients and mechanical effects limit the number of proliferating individuals to a small region close to the colony perimeter called the growth layer [21,22]. For mutations occurring inside the growth layer, most mutant offspring are concentrated in a relatively small number of enormously successful lineages that manage to remain at the front and “surf” on the expanding population wave [12]. As a consequence, clones of spontaneous neutral mutations often reach much larger sizes [16], and existing beneficial variants can sweep to high frequency much faster in microbial colonies than in well-mixed populations [21]. Conversely, deleterious mutations are predicted to remain at the population frontier for extended periods because genetic drift is strong at the front [8,19,36,43,47]. The quantitative outcome of the competition of selection and genetic drift in microbial colonies is determined by the local shape and roughness of the front [14,21], which in turn is determined by microscopic details, such as cell-cell adhesion or cell shape affecting the mechanical interactions between cells [18,30,31], although the direct mapping is typically unknown [14].

The evolutionary effects of fluctuations at expanding microbial population fronts have been studied in depth, but these studies have focused only on fluctuations associated with the growth, division, and random motion of cells, whose strength may depend on *intrinsic* properties of the microbial species, in homogeneous environments. However, any realistic microbial range expansion will experience varying degrees of environmental heterogeneity in the form of, e.g., nutrient or temperature gradients, or imperfections in the surface the population grows on. However, the effects of such environmental heterogeneity, which can be viewed as a source of *extrinsic* noise, on evolutionary dynamics in microbial populations have received much less attention. Efforts have concentrated mostly on simple temporal and spatial gradients in antibiotic concentration, which have been shown to facilitate the emergence of resistance in shaken cultures [37], microfluidic devices [51] and on agar plates [4], as predicted by theory [20,23,25–27].

The effects of spatial heterogeneity on evolutionary dynamics in expanding microbial populations has been studied in experiments only with neutral alleles in fixed geometries, such as isolated obstacles creating “geometry-enhanced” genetic drift [5,39]. These studies have shown that obstacles obstructing locally the advance of the front can doom lineages that happen to lie on the blocked part of the expanding population, whereas unobstructed lineages close to the edge of the obstacles obtain a boost as they fill the vacant area behind the obstacle. Even for neutral alleles, however, not much is known about the evolutionary dynamics in more complex heterogeneous environments. Moreover, selection during range expansions can dramatically alter the population structure over just a few generations [21], but how the action of selection is affected by environmental heterogeneity has so far remained completely unexplored.

Here, we study the impact of complex environmental disorder on the growth and evolutionary dynamics of microbial colonies. To this end, we introduce plasmid loss in *E. coli* as a model system for spontaneous mutations with tunable growth rate effects whose clones can be tracked under the microscope. By growing colonies on solid substrates with a weakly patterned surface with random microscopic features much bigger than individual cells, but much smaller than the whole colony, we find that environmental heterogeneity can overpower selection such that even strongly beneficial mutations are unable to establish at rates higher than expected for neutral mutations. Using a minimal computer model of populations expanding in randomly disordered environments, we show that dramatic changes in the efficacy of selection can arise from small changes in the degree of environmental heterogeneity. The limited efficacy of selection is a general consequence of a local “pinning” of the expansion front, whereby stretches of the front are slowed down on a length scale that depends on the structure of the environmental heterogeneity. This pinning focuses the range expansion into a small number of individuals with access to expansion paths, increasing the importance of chance and thus limiting the efficacy of selection. Our results may thus generalize to other spatially growing populations, such as biofilms, tumors, and invasive species, where the growing population front may be transiently hindered by the local environment.

## Results

### Experiments

We grew colonies from single cells of a strain of *E. coli* MG1655 on two different substrates: standard, “smooth”, agar plates as well as randomly patterned substrates (see Methods). The strain carries a plasmid that is costly to produce, resulting in a 20% growth rate disadvantage in plasmid-bearing cells compared to their plasmid-less (but otherwise isogenic) conspecifics. This strain loses the plasmid stochastically at a rate of about 5 × 10^-3^ per cell division (approximately independent of antibiotic concentration, see SI). The plasmid codes for a fluorescence gene and confers resistance to the antibiotic doxycycline (a tetracycline analog) such that by varying the amount of doxycycline in the growth media, the relative growth rates of the plasmid-bearing (“wild type”) and non-bearing (“mutant”) cells can be finely tuned from +20% to –15% (see SI Fig. S1). This allowed us to treat plasmid loss effectively as a spontaneous beneficial, neutral or deleterious mutation whose dynamics can be observed under the microscope. Our approach thus extends previous experimental model systems for evolutionary dynamics during microbial range expansion that employed either an initial mixture of wild-type and mutant cells [21,24,30,34] or were confined to spontaneous neutral [16] or deleterious [36] mutations. The ability to track spontaneous mutations in colonies grown from single cells is essential to ensure identical starting conditions in our experiments, allowing a quantitative comparison of the evolutionary outcomes between the two growth conditions. The primary readout of our experiments is the frequency *f*_MT_ (a proxy for the rate of adaptation of the population) and surviving number of mutant clones.

#### Homogeneous environment

Advantageous mutants increased in frequency *f*_*MT*_ rapidly as the colony grew: for *s* = 0.2, mutants made up roughly half of the total population (Fig. 2b) after 72 hours. Grown at higher antibiotic concentration, mutants became first neutral and eventually deleterious (for [*DOX*] *>* 0.3*µg/*ml). In such conditions, the mutant frequency decreased approximately exponentially with the fitness cost (SI Fig. S4) such that mutants made up only a small fraction of the final population.

**Figure 1.**
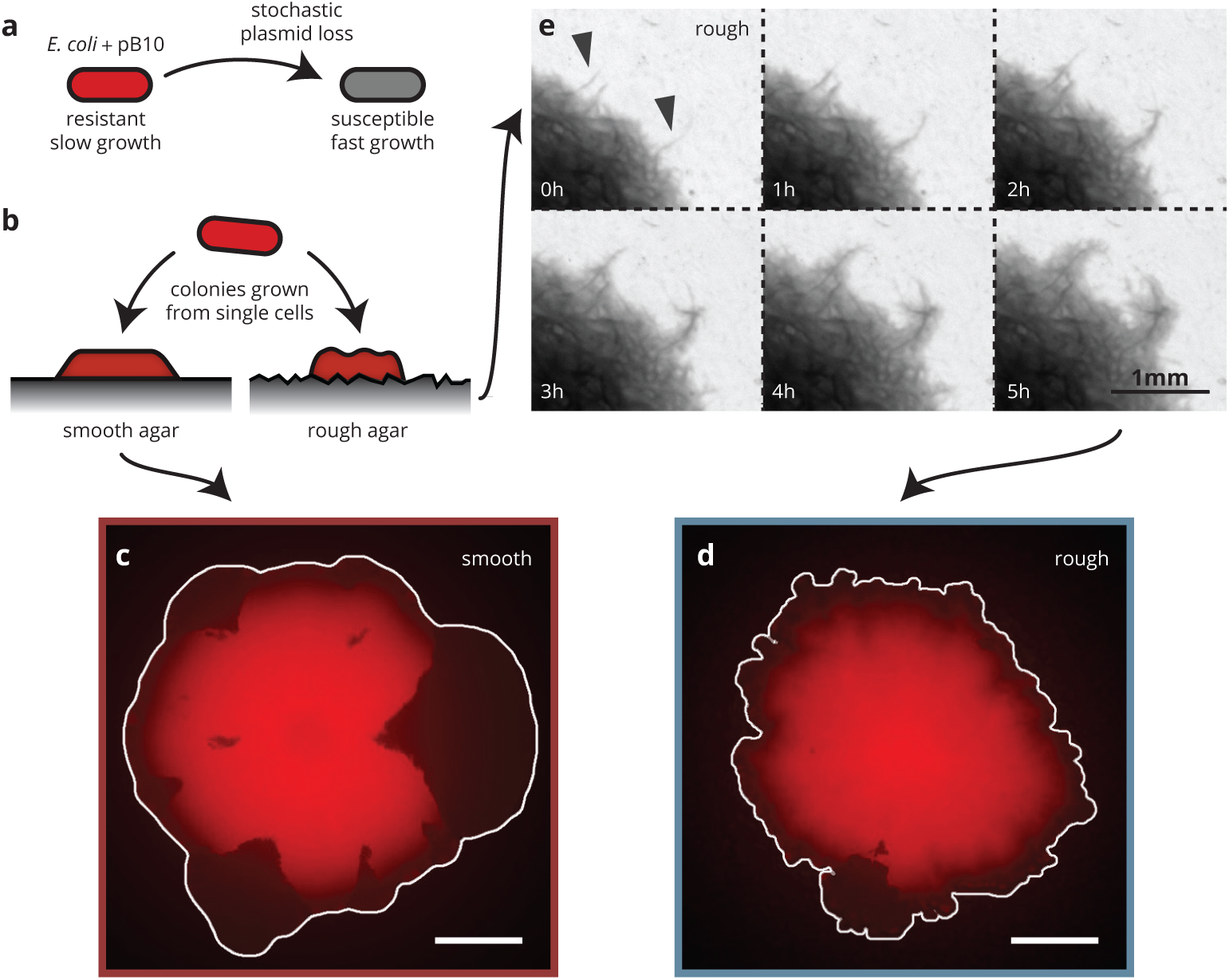
Substrates roughness changes bacterial colony morphology. (a&b) Colonies of *E. coli* were grown from single cells harboring a plasmid containing a fluorescence gene and a resistance cassette. Loss of the plasmid leads to non-fluorescent (“mutant”) sectors in the population that expand at the expense of the fluorescent (“wild-type)” population. Colony morphology depends on whether the colony grows on smooth (c) or rough (d, created by patterning the agar with filter paper) agar surfaces (Methods). (e) On rough surfaces, troughs in the surface direct growth along them, leading to locally accelerated regions that slowly widen and connect with the bulk of the population.

**Figure 2.**
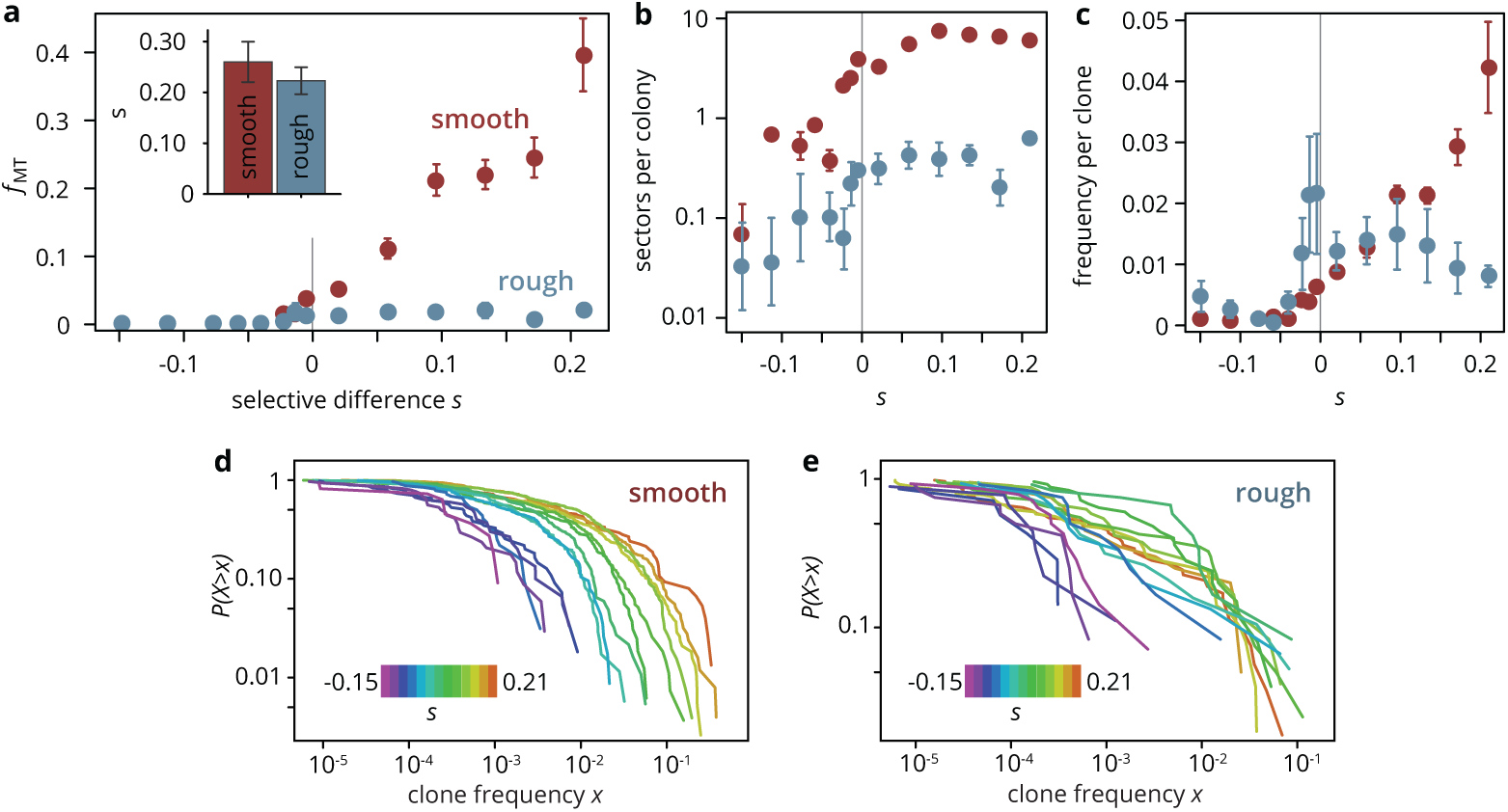
Substrate roughness limits the efficacy of selection in bacterial colonies. (a) The final frequency *f*_MT_ after 3 days of growth for mutants of a given fitness (dis)advantage *s* differed strongly depending on the roughness of the surface they were grown on, despite relative growth rate differences *s* being comparable (inset). On smooth plates, the mutant frequency *f*_MT_ increased with the selective advantage *s* of the mutants, whereas *f*_MT_ was independent of *s* for *s >* 0 on rough surfaces, and overall much smaller than in smooth colonies. The number of establishing sectors (b) was much smaller in rough colonies, and the average clone size (c) did not vary as strongly with *s* as in smooth colonies. (d,e) Clone size distribution *P* (*X > x*) for colonies grown on smooth and rough substrates. Deleterious mutants are shown in magenta tones; their clones are typically small. Neutral clones are shown green; their size distribution had a broad tail. Advantageous mutations (red tones) in smooth colonies were even more broadly distributed, as large sectors establish more often. By contrast, beneficial clones in rough colonies had size distributions that were indistinguishable from the distribution for neutral mutations.

The evolutionary success probability of individual mutations, measured by the establishment probability *u* to form a sector, also depended strongly the selective advantage (Fig. 2b), but was very low throughout. Even for the most advantageous mutants, we estimate *u* ∼ 10^-7^ per mutation (see SI Fig. S3), making sector formation an extremely rare event. The low success probability is a consequence of two processes: firstly, the mutation must occur in a favorable location, namely in the first layer of cells at the front of the population [21], which reduces the number of mutations eligible for sector formation by a factor of about 1000 (see SI). We estimate that about 2000 mutations per colony arose in favorable positions, each of which had an establishment probability of about 10^-3^. Secondly, each eligible mutation has to survive genetic drift, which in microbial colonies is manifest in the random fluctuations in the sector boundaries as a consequence of stochastic cell growth and division, and subsequent cell motion due to mechanical pushing of cells on each other [14,24,30].

Thus, most mutations will not manage to create sectors, but instead form ‘bubbles’, individual mutant clones that have lost contact with the front, the size of which we can extract from colony micrographs. The resulting clone size distribution *P* (*X > x*) (shown in Fig. 2d, e) is related to the site frequency spectrum in population genetics, where it can be used to predict rare evolutionary outcomes such as fitness valley crossing [49] and evolutionary rescue [16], and is well understood for toy models of microbial colonies [16,42]. For neutral mutations, the clone size distribution is expected to be broad up to a shoulder indicating the typical size of the largest expected bubble. In our experiments, we indeed observed a broad shoulder-like distribution for neutral mutations, consistent with earlier experiments using population sequencing [16] (Fig. 2e). For beneficial mutations, the larger number of bulging sectors created an even broader distribution with maximum clone sizes of almost half the population, while the distribution for strongly deleterious mutations was cut off at small clone sizes. The clone size distribution is consistent with our initial observation that a larger selective advantage *s* gave rise to a larger overall mutant frequency, but it also shows that even at the largest *s* ≈ 0.2, most mutant clones remained small, with more than half of the visible clones reaching frequencies of at most 1%.

#### Heterogeneous environment

To investigate the effect of environmental heterogeneity on colony growth and adaptation dynamics, we deposited filter paper with a pore size in the range of 9 to 20*µ*m onto melted agar and removed it after cooling and drying, creating an agar surface with random microscopic features. Colonies grown on these rough substrates (hereafter called “rough” colonies) had a rougher front line (Fig. 1d, SI Fig. S5) than those grown on smooth substrates (“smooth” colonies). The filter paper left grooves in the agar surface that the bacteria tended to colonize faster than the surroundings, leading the branch-like outgrowths that grew far ahead of the rest of the population and broadened as they were incorporated into the bulk of the colony (Fig. 1e).

Given the importance of the front morphology for evolutionary dynamics [14,21], we hypothesized that by changing the growth patterns of rough colonies, the structured agar surface should also impact the dynamics of spontaneous mutations. Indeed, the final mutant frequency *f*_MT_ in rough colonies was markedly different from what we found in smooth colonies (Fig. 2a, blue): while the neutral mutant frequency was roughly the same in both rough and smooth colonies, rough colonies showed no significant increase in mutant frequency with the fitness advantage *s* of the mutants, in contrast to smooth colonies, where the mutant frequency increased by a factor of 10 at the largest selective advantage *s* ≈ 0.2 compared to the neutral case. This effect did not stem from an altogether elimination of selection, however, as colonies of mutant and wild type grown separately on rough substrates showed a mutant fitness advantage (as measured by the radial growth rate) over the wild type consistent with the advantage they enjoyed on smooth substrates (Fig. 2a, inset).

The insensitivity of *f*_MT_ to selective differences can be broken down into a combination of two factors: the number of sectors was lower in rough colonies than in smooth colonies and constant for positive *s* (Fig. 2b), and the frequency per clone changed only slightly with *s* for *s >* 0 in rough colonies, whereas it increased approximately exponentially with *s* in smooth colonies (Fig. 2c). The discrepancy between mutant dynamics in smooth and rough colonies held also at the level of individual clones (Fig. 2d, e): for all *s >* 0, the clone size distributions obtained from rough colonies were virtually indistinguishable. By contrast, negative selection tended to decrease the size of mutant clones about equally in both smooth and rough colonies. Thus, beneficial clones in rough colonies behaved effectively neutrally, whereas deleterious mutations were fully affected by their growth rate disadvantage.

In summary, a microscopically randomly patterned growth surface had several effects on evolutionary dynamics in our colonies: it decreased the overall dependence of the final mutant frequency (or, equivalently, the rate of adaptation) on the selective effect of the mutation, it reduced the establishment probability of beneficial mutations, and it altered the distribution of clone sizes. These effects are large, despite the fact that the perturbation we impose is relatively weak. After all, the rough substrate is only distinguished from the smooth substrate by troughs and elevations much smaller than the whole population, or even a single beneficial mutant clone on smooth substrates, and colony growth rate differences are consistent for both substrate types (Fig. 2a, inset).

### Minimal model

How can a small change in environmental conditions have such a drastic effect on the evolutionary dynamics? To try to answer this question, we set up a minimal model of range expansions in the presence of environmental disorder to explore under which conditions small amounts of added disorder can generate the observed dramatic reduction in the efficacy of selection. Our model is based on the classical Eden lattice model [10] that is commonly used to model growing microbial colonies [6,16,21]. It has minimal ingredients: only cells with empty neighbors can divide, and a wild type can mutate upon cell division with probability *µ* to the mutant type carrying a fitness advantage or disadvantage *s*. Disorder sites (density *ρ*) confer a reduced growth rate *k* (0 *≤ k <* 1) to any individual growing on it. For *k* = 0, the disorder sites are impassable *obstacles*. The simplicity of the model allows us to explore exhaustively the whole parameter space in *k* and *ρ*.

The radial expansion speed of the colonies depends on both the obstacle density *ρ* and the growth reduction factor *k* (Fig. 3b). For relatively weak growth rate reduction at obstacles sites (i.e., *k* ≈ 1), the expansion speed decreases slightly with the obstacle density. For increasing growth reduction, i.e., for *k* → 0, the expansion speed decreases first slowly and then rapidly as the density reaches a critical value *ρ*_*c*_ ≈ 0.4 (dotted line). For impassable obstacles (*k* = 0) at densities *ρ > ρ*_*c*_, the obstacles form a closed ring around the incipient colony and prevented further growth (Fig. 3b, black line); thus, there is a phase transition at a critical density *ρ*_*c*_ ≈ 0.4. This transition is called the pinning transition of the interface (discussed in detail in the SI) and corresponds to the scenario where the colony can no longer percolate through the network of obstacles, suggesting that *ρ*_*c*_ is equivalent to the site percolation threshold 1 – 0.592… = 0.407… [3,7]. Close to the phase transition, small changes in obstacle density can have dramatic effects: not only does the colony expansion speed decrease rapidly, but as we shall see below, it also impacts evolutionary dynamics, such as the establishment probability of beneficial mutations and the final frequency of mutants.

**Figure 3.**
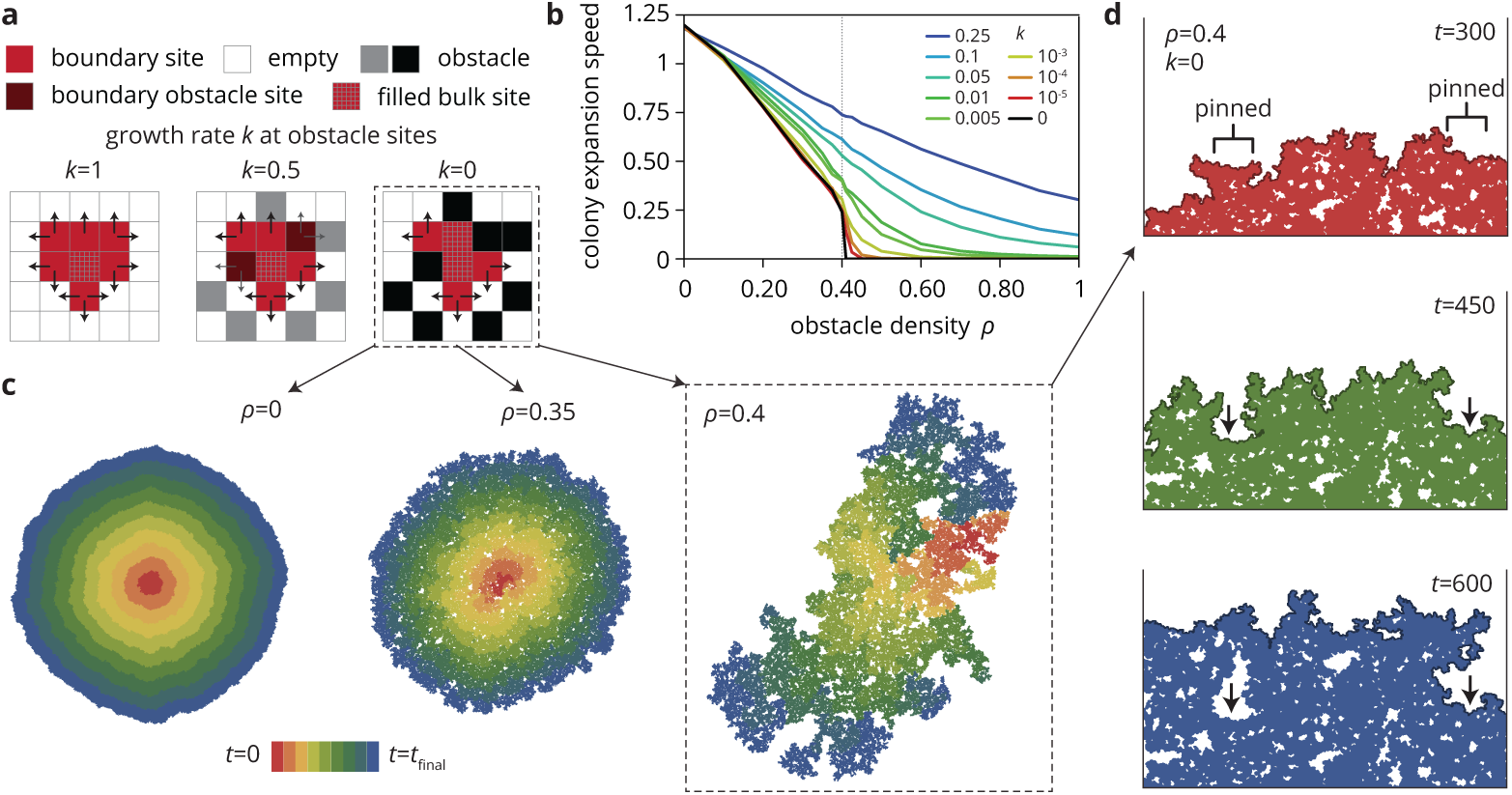
A minimal model captures the change in colony morphology in disordered environments. (a) The simulation proceeds by allowing cells with empty neighbors (“boundary sites”) to divide into empty space. Disorder sites (gray/black, density *ρ*) are randomly distributed over the lattice. (b) As the growth rate *k* at disorder sites is reduced, the colony expansion speed decreases as a function of their density. For impassable disorder sites (“obstacles”, *k* = 0), the expansion speed vanishes above a critical obstacle density *ρ*_*c*_ ≈ 0.4, beyond which the population was trapped and could not grow to full size. (c) As *ρ* approached *ρ*_*c*_ colony morphology became increasing fractured. Colors from red to blue show the colony shape at different times. (d) Near the critical obstacle density *ρ*_*c*_ ≈ 0.4, only few individual sites near the front have empty neighbors, leading to local *pinning* of the front (arrows), where the front can only progress by lateral growth into the pinned region.

In the presence of high obstacle densities, the interface is characterized by a characteristic length scale that diverges as the critical density *ρ*_*c*_ is approached. Over this length scale, the interface is *pinned*, i.e., it cannot advance locally. This pinning effect, which we also observed in our experiments (see Fig. 2e), is indicated as arrows in Fig. 3d. As a result of the local pinning of the colony interface, the colony morphology depends on the density of obstacles, most drastically for impassable obstacles on which we concentrate in Fig. 3c. Without obstacles, the colonies are compact and relatively smooth. At inter-mediate obstacles densities, colonies are punctured by small holes and the overall density of the colony decreases. At the critical density *ρ*_*c*_ the colony is characterized by the fragmented morphology of percolation clusters with a large number of holes and a very rough exterior (see SI Fig. S7 for a quantitative analysis of the colony interfaces). In the following, we investigate how this change in colony morphology affects the evolutionary dynamics.

We begin by replicating the experimental situation to assess the efficacy of selection in the presence of environmental heterogeneity. We simulated mutations conferring a selective advantage *s* (i.e., increasing the growth rate by a factor 1 + *s*), shown in Fig. 4. Relatively weak obstacles (growth rate *k* at obstacle site *k* = 0.1, Fig. 4a) only have a mild effect on the mutant frequency *f*_MT_. As in our experiments, *f*_MT_ increases roughly exponentially with *s*, albeit slightly weaker at intermediate *ρ* ≈ 0.5 than at the extremes *ρ* ≈0 or *ρ* ≈1. This reduction in the sensitivity of *f*_MT_ to *s* becomes much more dramatic as the growth rate on disorder sites is decreased (see Fig. 4b for the extreme case *k* = 0). In addition, as the critical density *ρ*_*c*_ is approached, the frequency of neutral mutants increases, resulting in mutant frequencies that are elevated relative to the homogeneous scenario for neutral and deleterious mutations, but reduced for beneficial mutations.

**Figure 4.**
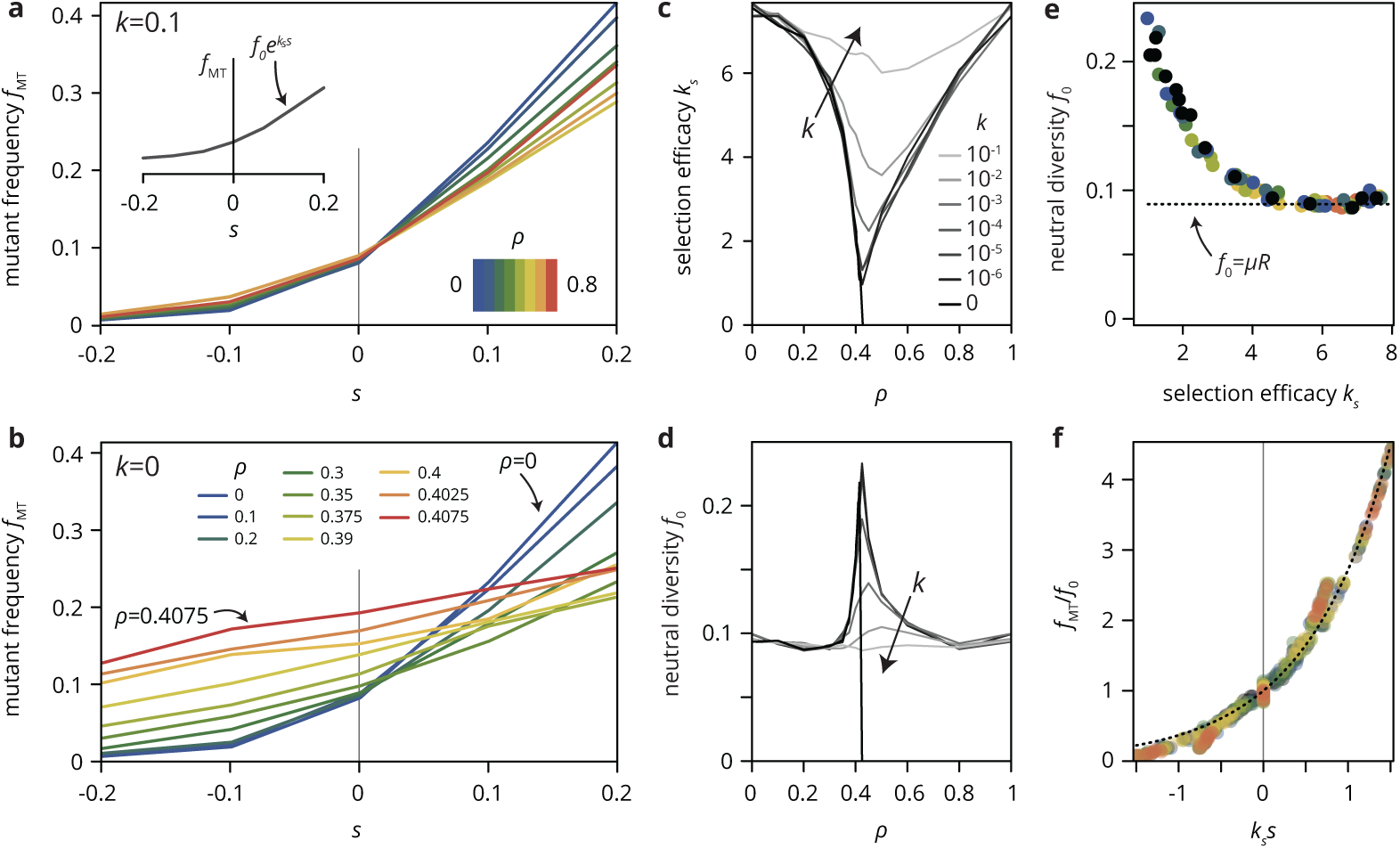
Consequences of simulated environmental disorder for the evolutionary dynamics. (a,b) The mutant frequency *f*_MT_ as a function of the selective advantage *s* of the mutants always increases with *s*, but it depends on the density *ρ* of and the growth rate *k* on disorder sites. (c,d) Parametrizing the curves for various values of *k* and *ρ* with 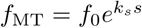 (see panel a, inset), we fit the neutral diversity *f*_0_ (shown in panel d) and the selection efficacy *k*_*s*_ (panel c), which serves as an inverse selection scale, describing the dependence of *f*_MT_ on *s*. (e) A scatter plot of the fit parameters *f*_0_ and *k*_*s*_ for all values of *k* and *ρ* reveals that they are not independent parameters, but that *f*_0_ entirely determines *k*_*s*_ and vice-versa (for a given population size *N* and mutation rate *µ*; here, *N* = 10^5^ and *µ* = 0.0005). The dashed line represents the expect neutral diversity for a circular colony, where *f*_0_ = *µR*, with 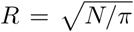 the colony radius [16]. (f) Rescaling all curves by fitted values of *k*_*s*_ and *f*_0_, all points fall onto a master curve given by 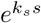 (dashed line), justifying the parametrization introduced in (a).

To quantify the effects of varying *k* and *ρ* and summarize the simulation results over many parameter combinations, we introduce the selection efficacy *k*_*s*_ by parametrizing the mutant frequency with an exponential function 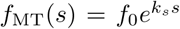. This choice is merely heuristic, but is justified by rescaling mutant frequency curves for a range of values of *ρ* and *k* by the fitted values for the neutral diversity *f*_0_ and the selection efficacy *k*_*s*_, which collapses all data onto a single master curve given by a simple exponential (Fig. 4f). The selection efficacy *k*_*s*_ has a minimum near *ρ*_*c*_, which is increasingly sharp for decreasing *k*. It vanishes entirely as *ρ* approaches *ρ*_*c*_ for obstacles (*k* = 0), indicating that selection is completely unable to affect the final mutation frequency in this limit. The virtual independence of the evolutionary dynamics of the per-capita fitness *s* holds even at the scale of individual clones, whose size distributions for different values of *s* are practically indistinguishable for obstacle densities near *ρ*_*c*_ (SI Fig. S6). Thus, while we find a proper phase transition only for *k* = 0, the percolation transition is also manifest in the evolutionary dynamics in populations grown in generic heterogeneous environments. As a consequence, tiny changes in environmental parameters near a non-trivial critical obstacle density *ρ*_*c*_ can have a dramatic effect on colony morphologies and evolutionary dynamics. The direct connection between colony morphology and evolutionary dynamics is underscored by the dsicovery that the two descriptive parameters, the selection efficacy *k*_*s*_ and the neutral diversity *f*_0_, introduced as independent parameters measured directly from the simulations, are not independent in practice (Fig. 4e). Plotting *k*_*s*_ vs. *f*_0_ for various choices of *k* and *ρ* reveals that the two parameters represent two sides of the same coin: environmental disorder alters the growth pattern of the colony, which in turn affects both the neutral diversity and the selection efficacy.

From a classical population genetics perspective, the fact that the addition of extrinsic noise effectively weakens selection is not surprising, as other sources of noise, such as small population sizes, are known to push evolutionary dynamics towards the neutral limit [17]. However, the environmental heterogeneity in our simulations changes the evolutionary dynamics on a fundamental level that is not consistent with a mere increase in total noise. To show this, consider the neutral diversity *f*_0_ in Fig. 4e, which corresponds to the rate at which neutral mutations accumulate in the population. On average, this rate is *µ*(*N/π*)^1/2^ since a fraction *µ* of cells at the population front acquire new neutral mutations in every generation, and the front scales as the square root of the population size *N* [16]. Importantly, this result in independent of the level of noise in the system since it concerns only the average over many populations. Since the population size is the same across all our simulation, we would expect the same neutral diversity for all parameter values *ρ* and *k*, but our simulations show clear systematic deviations from the expected value (dotted line in Fig. 4e); in particular, for small *k*, the neutral diversity *f*_0_ has a pronounced maximum near *ρ*_*c*_ (Fig. 4d).

To further characterize the qualitative changes on the neutral dynamics induced by environmental heterogeneity, we computed the spatially resolved phylogenetic tree of the population, obtained by tracing the lineages of all individuals at the population front back to the origin. For simplicity, we concentrate on the case of no (*ρ* = 0) and critical (*ρ* = *ρ*_*c*_) obstacles (i.e., *k* = 0). As shown in Fig. 5a, b, the tree has a vastly different appearance depending on the environmental heterogeneity. Without obstacles (panel a) the lineages are relatively straight and roughly aligned with the radial direction. By contrast, at the critical obstacle density, where the colony has a rough exterior, lineages are erratic and often have segments oriented perpendicular to the radial direction. The lineages shown in Fig. 5a,b are intimately related to the shape and orientation of individual neutral clones (Fig. 5g-j). Mutant clones have an approximately ellipsoidal shape oriented preferentially along the radial direction in the absence of heterogeneity, whereas they have essentially random orientations in rough colonies (Fig. 5i), in agreement with the observation that lineages lose their radial orientation as the number of obstacles increased. Similarly, the scaling of the clone width 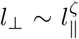 with its length *l*_*||*_ changed from *ζ* = 2/3 for *ρ* = 0 (consistent with KPZ interface statistics [16]), to *ζ* ≈ 0.95 for *ρ* = *ρ*_*c*_ (Fig. 5j), indicating roughly isotropic neutral clones.

**Figure 5.**
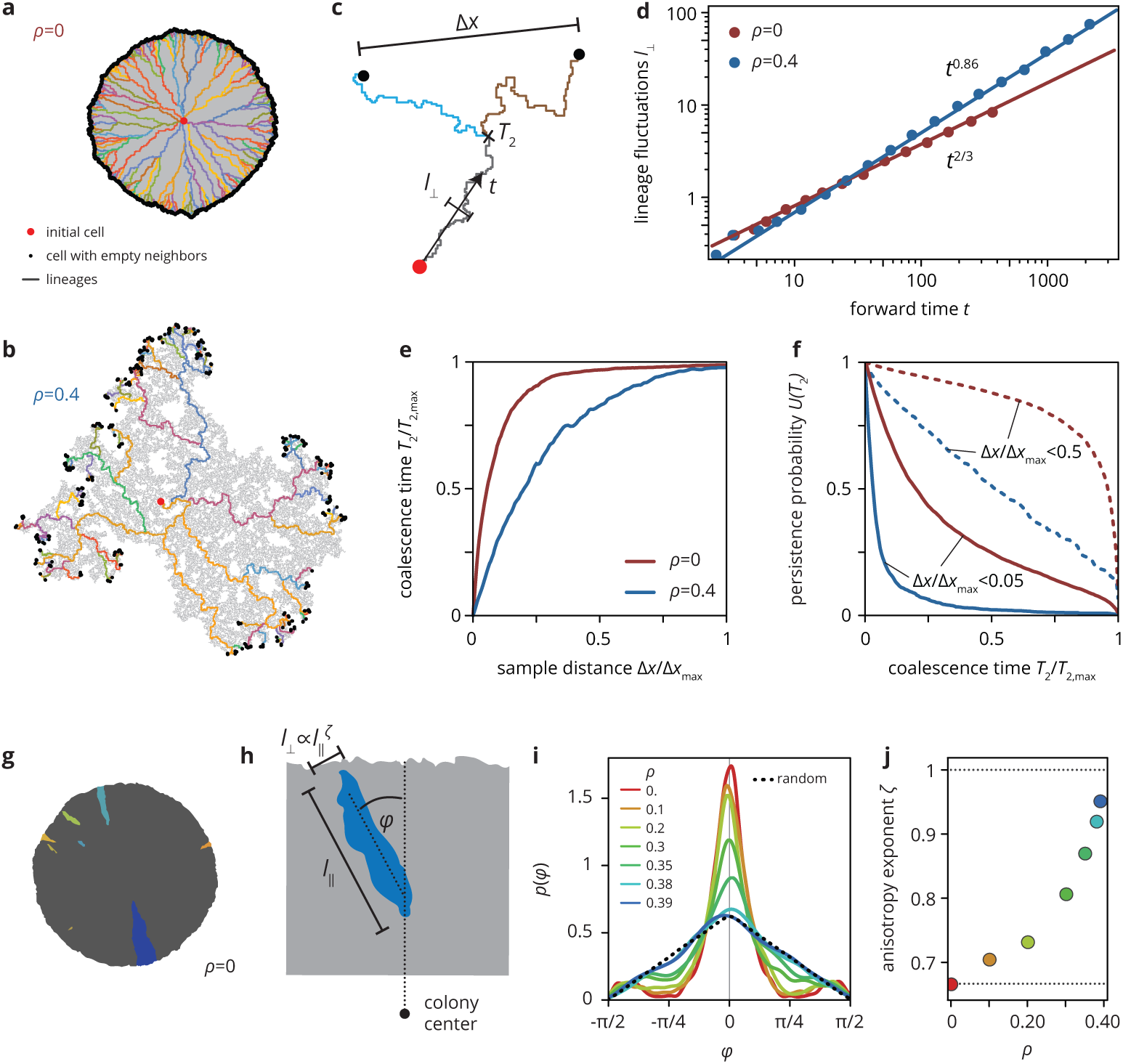
Neutral lineage dynamics change with environmental heterogeneity. Lineage trees extracted from simulated colonies without (a) and with (b) environmental heterogeneity. (c) The lineage structure is characterized by the fluctuations *l*_⊥_ of the lineages in forward time *t*, and the pair coalescence time *T*_2_ (backwards in time) of two samples that are a distance Δ*x* apart. (d) The lineage fluctuations depend on the environmental disorder: without disorder, lineage fluctuations scale as *t*^2^/^3^, whereas the lineages are rougher in strong disorder, scaling as *t*^0^.^86^. Strong lineage fluctuations in heterogeneous environments also impact the mean pair coalescence time of two samples a distance Δ*x* apart by allowing lineages from distant regions of the colony to coalesce earlier than without environmental disorder (panel e). As a consequence, the persistence probability *U* (*T*_2_), i.e., the probability that two sampled lineages have a pair coalescence time greater than *T*_2_ is typically greater in homogeneous than in heterogeneous environments, even when conditioning on small distance between samples (solid lines, Δ*x <* 0.05Δ*x*_max_; dashed lines, Δ*x <* 0.05Δ*x*_max_).

To quantify how environmental heterogeneity impacts neutral lineage dynamics, we focused on the strength of lineage fluctuations and the coalescence “time” measured in lattice sites (see Fig. 5c). First, the lateral lineage fluctuations *l*_⊥_∼ *t*^*ξ*^ as a function of distance *t* from the origin are not only rougher in absolute value, but also in terms of their scaling in rough colonies. Whereas in the standard Eden model we recover the known scaling *ξ* = 0.66 ± 0.006 [29], we find a larger scaling exponent *ξ* = 0.86 ± 0.006 in rough colonies (*ρ* = 0.4). This is consistent with the corresponding change of the statistical properties of the colony interface, which transitions from the Kardar-Parisi-Zhang (KPZ) universality class to the quenched Edwards-Wilkinson (QEW) universality class (see SI). Second, the increased roughness of lineages is also reflected in the number of successful lineages emanating from the initial population founder. We quantify this by computing the pairwise coalescence time *T*_2_ (Fig. 5e,f), i.e., how long ago two individuals a distance Δ*x* apart at the front had their most recent common ancestor. We find that, for a given sample distance, the relative coalescence time and persistence probability (i.e., the probability of not having a common ancestor until time *T*_2_, panel f) is always smaller in the presence of obstacles. This indicates that fewer lineages reach the population edge in the presence of environmental heterogeneity. This makes intuitive sense from the phylogenetic trees shown in Fig. 5a, b, where in rough colonies all individuals at the front coalesce quickly into a small number of large lineages. In summary, the observed differences in selection efficacy, neutral diversity, lineage scaling properties, and (neutral) coalescence structure between populations expanding in homogeneous or heterogeneous environments rule out the interpretation of environmental disorder as simply another additive noise. Instead, we find that environmental disorder can fundamental alter the population genetics of range expansions.

## Discussion

Evolution can be viewed as the result of a competition between the deterministic force of selection and various sources of randomness: firstly, intrinsic noise, which encompasses genetic drift, the nature and timing of mutations, etc., only depends on the inherent properties of the population, such as the species (including its microscopic characteristics like adhesion strength, cell shape, or growth rate) and the population size. The second source of randomness is extrinsic noise, by which we mean spatio-temporal gradients and fluctuations in environmental conditions, such as temperature, nutrient availability, or antibiotic concentration. To fully understand evolutionary dynamics, the relative strength of all three factors have to be taken into account. In evolution experiments, extrinsic noise is typically either deliberately prevented, or added in a very controlled fashion, such as periodic changes in conditions.

Here, we have shown that extrinsic noise in the form of random environmental disorder can dramatically impact the fates of spontaneous mutations in microbial colonies. By leveraging plasmid loss in *E. coli* as a model system for spontaneous mutations with tunable growth rate effect, we have overcome previous experimental limitations, which required the mixture of two strains to study the effects of beneficial mutation. We have confirmed previous theoretical predictions about the evolutionary dynamics of spontaneous mutations in microbial colonies grown on homogeneous substrates: mutants with larger selective advantages are more likely to establish clonal sectors that expand rapidly and quickly become highly abundant in the population; overall, however, even very advantageous mutations are undividually extremely unlikely to be successful, leaving the evolutionary fate of the population in the hands of a few lucky clonal lineages.

By growing colonies on heterogeneous substrates, we found that microscale ridges and troughs in the growth substrate were enough to reduce the ability of beneficial mutations to establish and expand. Our minimal model simulations showed that this reduction in selection efficacy on heterogeneous substrate can be explained by a local pinning of the colony front. Since mutations occur only within the growing population at the front, the properties of the front dictate the evolutionary dynamics, including the strength of selection and the size of individual clones. Local pinning impacts the dynamics at the front by reducing the expansion speed of some parts of the population, leading to an effective reduction in the number of expansion paths that can actively contribute successful mutations. Thus, only a few lucky lineages will be able to find the paths along unpinned front positions; most lineages will get stuck in dead-ends. Given local establishment, lineage success is then roughly independent of the fitness of the mutants; whether a sector can form or not depends *entirely* on where the mutation arises and not at all on its growth rate. Similarly, the size of mutant clones is constrained by the network of obstacles, and whether and when a given mutant clone goes extinct depends only marginally on the fitness of its founder. In this sense locally pinned expansions bear little resemblance to unconstrained radial expansions (see Fig. 6). Rather, expansions along each available path more closely correspond to linear expansions, e.g., along a coastline, where mutations spread deterministically after local establishment [15]. Overall, the observed differences in selection efficacy, neutral diversity, lineage scaling properties, and (neutral) coalescence structure between populations expanding in homogeneous or heterogeneous environments rule out the interpretation of environmental disorder as simply another additive noise. Instead, we find that environmental disorder can alter the population genetics of range expansions at a fundamental level.

**Figure 6.**
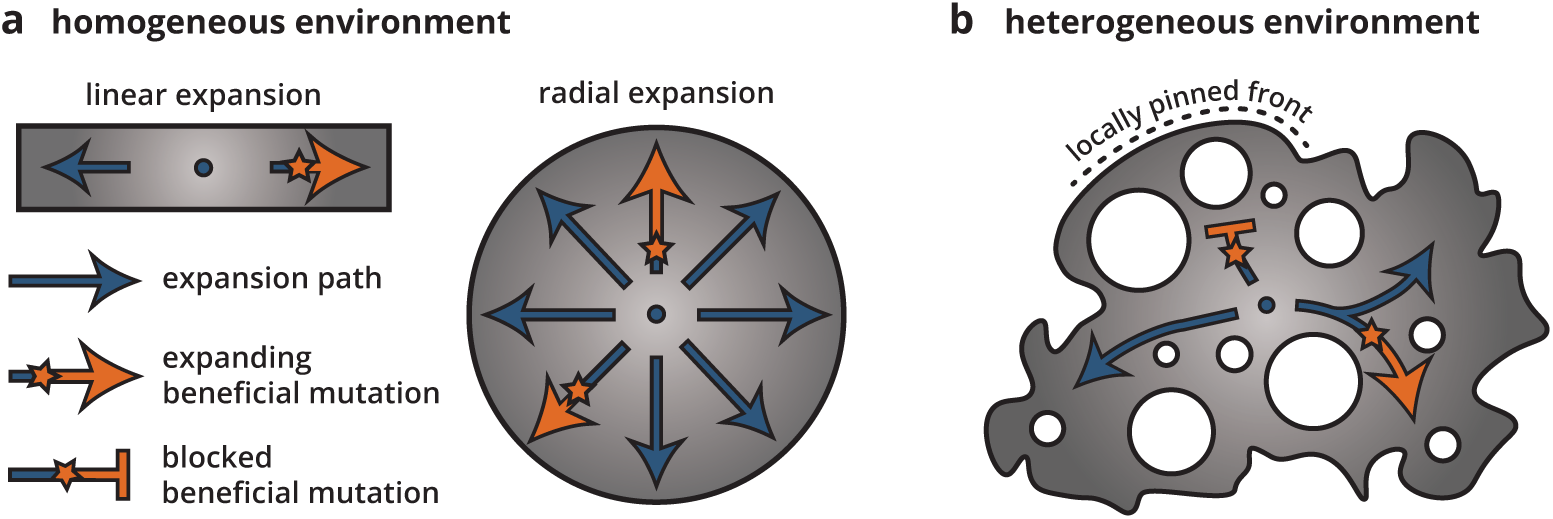
Cartoon of how environmental heterogeneity shapes evolutionary dynamics during range expansions. In homogeneous environments (a), a locally established beneficial mutation (orange star), having survived genetic drift, can expand freely (arrows). By contrast, in heterogeneous environments (b), a beneficial mutation can become trapped in pinned stretches of the population, whereas expansion along open paths resembles a one-dimensional expansion.

Environmental disorder arguably impacts not only the rate of adaptation due to beneficial mutations. Since deleterious mutations are typically more numerous than beneficial ones, environmental disorder may also increase the chances of an overall *decrease* in population fitness through the accumulation of deleterious mutations, which is already more likely in range expansions than in well-mixed populations [19,36,43]. Thus, heterogeneities in the environment may not only slow down the process of adaptation but also lead to entirely different long-term evolutionary outcomes. Since many mutations conferring resistance to antibiotics are often associated with a fitness cost, environmental disorder may also favor the evolution of resistance in microbial populations in this way. However, our results on the fates of deleterious mutations remain ambiguous: while our simulations predict that deleterious mutations should be more successful in disordered environments, our experiments found no significant differences in the size of deleterious clones between smooth and rough colonies. A potential reason for this discrepancy is may be that the disorder imposed in our experiments is not correlation-free. A beneficial mutation has to overcome genetic drift, and to do so, it must grow to a lateral size large enough for selection to take over [21]. However, if the characteristic length scale of the environmental disorder is smaller than this “establishment size”, then the evolutionary dynamics is effectively neutral. On the other hand, a deleterious mutation born on a ridge or in a trough never grows to large enough size to “see” the disorder in the first place and thus its dynamics are largely unaffected by the environmental disorder.

While we have focused here on microbial populations, we expect our principal result of reduced selection efficacy in disordered environments to generalize to other dense cellular populations, such as tumors and biofilms, but also to macroscopic range expansions, as well. After all, when a population undergoes a range expansion, it will arguably not experience a completely homogeneous environments: at the very least, some areas will be more hospitable than others, but other parts of the environment may also be entirely inaccessible to the population because of, e.g., rivers and lakes, a strong local competitor or predator, or lack of resources. Environmental heterogeneity is thus arguably the rule rather than the exception. Our results suggest that since range expansions in strongly heterogeneous environments can generate approximately neutral patterns of genetic diversity from mutations carrying significant fitness effects, attempts to interpret such patterns in invasive species, or generally species having undergone recent range expansions, must take into account the role the environment plays in shaping these patterns.

## Acknowlegdments

The authors thank Jona Kayser, Diana Fusco, Jayson Paulose, and all members of the Hallatschek lab for helpful discussions. Research reported in this publication was supported by the National Institute of General Medical Sciences of the National Institutes of Health under award R01GM115851, a National Science Foundation CAREER Award (#1555330) and a Simons Investigator award from the Simons Foundation (#327934).

## Supplementary information

### Experimental methods

#### Strains and growth conditions

We used an *E. coli* MG1655 strain transformed with the plasmid pB10 [45]. pB10 is a 65kB plasmid isolated from sewage sludge that confers resistance to several antibiotic resistance including tetracyclines and has an inserted RFP gene. Hence, cells containing pB10 (“wild type”) are red fluorescent and resistant to tetracycline. The plasmid is lost sporadically [9], and the resulting cells (“mutants”) are non-fluorescent and susceptible to tetracycline, but display a higher growth rate in the absence of antibiotics (characterized below). We refer to the loss of the plasmid as a “mutation” of known fitness effect and occurrence rate, both of which we characterize below.

All experiments were performed in LB at 37°C in a humidified environment. For solid media, 2% agar was added before autoclaving. Varying concentrations of doxycycline, a tetracycline that displays higher stability in agar plates than tetracycline itself, were added to freshly autoclaved media after cooling to about 60°C and poured immediately. Plates were dried in the dark for at least 24h before use.

#### Fitness measurements

We measured the fitness difference *s* between wild type and mutant cells using the colliding colonies assay [21,34]. Briefly, a mutant clone was first isolated and then grown independently of the wild type overnight. In the wild type, the plasmid was maintained by adding 10*µ*g/ml doxycycline to the overnight culture. After growth overnight, cultures were diluted 1:10, grown for about 1.5h, and then washed twice in PBS to remove residual doxycycline. 1*µ*l droplet of each strain were spotted on agar plates about 2mm apart. After drying, colonies were grown for 3 days and then imaged under the a Zeiss Axiozoom v16 microscope. The resulting images were used to estimate fitness differences by fitting a circle onto the mutant-wild type interface. The results are shown in Fig. S1: without doxycyline, mutants have a 20-25% advantage over the wild type. Both strains have equal growth rate around ≈0.35ug/ml, and the mutants grow more slowly than the wild type at higher concentration of doxycycline.

For the growth rate measurements in Fig. 2, colonies were grown from single cells on both rough and smooth plates in a temperature-controlled growth chamber and imaged overnight on a Zeiss Axiozoom v16 microscope. The resulting time lapse movies were binarized and the colony areas extracted.

#### Mutation rate experiment

To measure the rate of plasmid loss (“mutation rate”), we grew 48 well-mixed populations from a small number of wild-type cells for about 7 generation (i.e., from about 10 to about 1000 cells). The inoculum did not contain any mutant because the culture used to inoculate the populations contained selective amounts of tetracycline. The populations were grown either without doxycycline or at 1*µ*g/ml doxycycline, which corresponds to the low and high end of concentrations used in our experiments, respectively. After 7 generations, each population was plated and the number of red (WT) and gray (MT) colonies was counted via automated image analysis. The resulting frequencies of mutants were used to infer the mutation rate by computing the maximum likelihood against simulations of the process at different mutation rates and fitness differences, as follows.

#### Statistical inference of mutation rate

To estimate the mutation rate, we performed maximum likelihood estimation based on probability density distributions obtained from simulations, as follows: starting from a Poisson distributed number of initial cells, 48 populations go through about 7 generations, where every wild type has a chance *µ* per division to produce a mutant. Mutant cells grow at a growth rate (1 + *s*) relative to the wild type. We performed 50000 simulations for each value of *s* and *µ* and computed the likelihood of each parameter combination *θ* = *s, µ* as

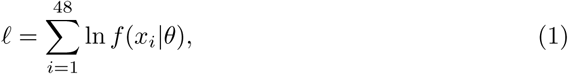

where *f* (*x*_*i*_) is the probability of observing *x*_*i*_ under the simulation model, which we estimated from the simulation histograms. We treat *s* as a free parameter that we can later compare to the experimentally measured value. The precise value of *s* does not affect the inferred value of *mu* very strongly. This is because the number of generations is small in our experiment and a faster-growing mutant can gain at most a factor of four more cells than the wild type. The global maximum likelihood value *µ\** is obtained for *s* = 0.3 and *s* = –0.05 for doxycycline concentrations of 0*µ*g/ml and 1*µ*g/ml, in good agreement with our measured values of *s* (see Fig. S2). The error is estimated from the curvature of the likelihood as 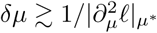. The results are *µ*_0*µ*g/l_ = 0.003 ± 0.00055 and *µ*_1*µ*g/l_ = 0.009 ± 0.00068.

#### Main experiment and analysis

Our main experiment consists in the growth of colonies from single wild-type cells on agar plates (each containing 20ml of LB + 2% agar) containing varying concentrations of doxycycline. The agar plates were either *smooth*, standard agar plates, or *rough*. Rough plates were created by pouring the agar at a temperature of about 60°C and then gently lowering filter paper (VWR Grade 410 Filter Paper, Qualitative, pore size between 9 and 20*µ*m) onto the liquid agar, where it remained until the agar had solidified. The filter paper was then removed from the hard agar surface with tweezers, resulting in a patterned agar surface.

Overnight culture of the wild type grown in LB with 10*µ*g/l doxycycline was washed and diluted in PBS to give between 3 and 10 colonies per plate. About *n* = 20 – 30 colonies per condition were analyzed (except for smooth plates without doxycycline, where *n* = 8). After 72 hours of growth, the colonies were imaged on a Zeiss Axiozoom v16 microscope and the resulting images binarized to create a mask of the colony. Mutant clones we found manually with ImageJ. The mutant frequency per colony was then measured as the total mutant area divided by the total area of the colony.

### Simulations

To simulate growing bacterial colonies, we employ an Eden model [10] on a square lattice that we generalized to include mutations with fitness effect *s* and environmental disorder. To simulate environmental disorder, we first initialize the lattice with a number of disorder sites at a density *ρ*. Disorder sites are characterized by a reduced growth rate *k* (0 *≤ k <* 1). For *k* = 0, the disorder sites are impassable (we call this type of disorder sites *obstacles*); equivalent models have also been used to simulate epidemics, where the obstacles represent resistant sites [28]. Without disorder, our model is identical to that used in Ref.s [16] and [21], and its interfaces are known to be well described by a standard model for stochastically growing interfaces, the KPZ equation [29].

The population is initiated with a single filled site in the center. In each time step, a site with empty neighbor sites is chosen with probability proportional to its growth rate *k* × (1 + *s*) to divide into a randomly chosen empty neighbor site (Fig. 3a). Upon division, a wild-type site acquires a single mutation with probability *µ*, potentially conferring a fitness advantage or disadvantage *s*; already mutated sites do not acquire further mutations. The populations are grown until the same number of lattice sites *N* is filled; for strong environmental noise this results in colonies that are optically larger (see, e.g., Fig. 3c).

### Theory

Interfaces created by Eden model simulations fall into the KPZ universality class, governed by the KPZ equation for the height *h*(*x, t*) [13,29]. In one dimension, starting from a line in a simulation box, the colony surface is described by

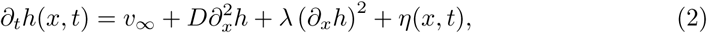

where *v*_*∞*_ is the final speed of front propagation and *η*(*x, t*) is zero-mean Gaussian random noise *δ*-correlated in space and time describing the noise associated with the growth process. This equation generates a set of characteristic exponents that govern the roughness of the colony front and of sector boundaries. In particular, the surface height is described in terms of its root mean squared fluctuations around the mean by a Family-Viscek scaling relation [48]

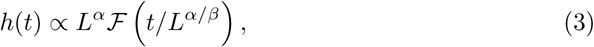

where

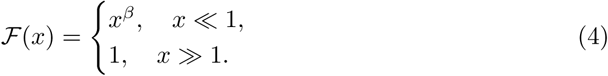

The KPZ universality class is characterized by the roughness exponent *α* = 1/2 and the temporal exponent *β* = 1/3; if *λ* = 0, the resulting universality class is called the Edwards-Wilkinson (EW) universality class characterized by *α* = 1/2 and *β* = 1/4. The ratio *z* = *α/β* is sometimes called the dynamical exponents. It relates the size of lateral fluctuations *l*_⊥_ to the time *t* as

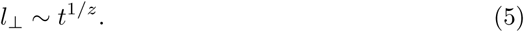

This relationship explains the fluctuations in sector boundaries in *E. coli* colonies and Eden model simulations, and can be used to derive exponents for the site frequency spectrum and establishment probabilities in Eden model colonies [16,21,24]. In the simulation presented here, the scenario without environmental disorder is described the eq. (2), explaining the site-frequency spectrum and the anisotropy exponents *ζ* = 2/3 = 1*/z* and the lineage fluctuations in Fig. 5.

The effect of environmental *quenched* disorder on the kinetic roughening of interfaces has been investigated in a range of experiments (see Ref. [3] for a review). Exponents obtained from experiments are in the range of *α* ≈0.6 … 0.9. To model driven interface growth in disordered media, quenched environmental disorder can be included by considering a noise term *ζ*(*x, h*(*x, t*)) that does not explicitly depend on time [32].

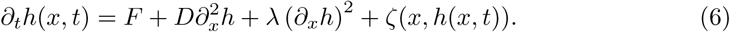

Here, *F* is driving “force” fulfilling the same role as *v*_*∞*_ in eq. (2). Since the noise explicitly depends on the interface position, *F* cannot be transformed away and thus emerges as a new parameter that can be thought of as a force pushing the interface through the disordered media. An important consequence of quenched noise is the emergence of a critical force *F*_*c*_ below which the interface becomes pinned [46]. For *F > F*_*c*_, a depinning transition takes place that is well characterized in 1+1 dimensions [1]. For 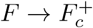, large regions of the interface of size *ξ* ∼*|F*_*c*_ *F|* ^*-ν*^ are pinned, and the front speed increases as |*F – F*_*c*_*|* ^*θ*^ (see Table 1).

**Table 1.** Characteristic exponents for the KPZ and QEW universality classes and measured in our simulations, which are in good agreement with the literature values in Ref. [1].

| Exponent | KPZ | This study ($\rho = 0$) | qEW | This study ($\rho = \rho_c$) |
| --- | --- | --- | --- | --- |
| $\alpha_{\text{loc}}$ | 1/2 | $0.5 \pm 0.05$ | $0.92 \pm 0.04$ | $0.9 \pm 0.05$ |
| $\alpha_G$ | 1/2 | $0.5 \pm 0.05$ | $1.23 \pm 0.04$ | $1.15 \pm 0.05$ |
| $\beta$ | 1/3 | $0.3 \pm 0.03$ | $0.86 \pm 0.03$ | $0.78 \pm 0.05$ |
| $\theta$ | N/A | N/A | $0.24 \pm 0.03$ | $0.21 \pm 0.03$ |
| $\zeta$ | 2/3 | $0.66 \pm 0.02$ | - | $0.86 \pm 0.04$ |

Simulations and numerical integrations of eq. (6) have characterized the pinned and moving phases and uncovered two universality classes as 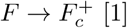 [1]: if *λ* diverges near the depinning transition, one speaks of the (quenched) QKPZ universality class; its exponents *α* = *β* ≈ 0.633 in the pinned phase are understood analytically through an analogy with the directed percolation class, whereas in the moving phase *α* ≈ *β* ≈ 0.75. If *λ* → 0 instead, one speaks of the quenched Edwards-Wilkinson (QEW) universality class with *α* ≈ 0.92 and *β* ≈ 0.82 in the moving phase (see also Table 1); a functional renormalization group calculation gives *α* = 1 and *ν* = 1/(2 *- α*) in one dimension [41]. For *F > F*_*c*_, there is a transition from the QKPZ/QEW universality class to the appropriate universality class with annealed noise.

Our generalized Eden model simulations with obstacles exhibits the same pinning transition for an obstacle density *ρ* ≈ 0.4. At the transition, the resulting colonies are (site) percolation clusters on the square lattice, from whose interfaces we measure exponents that are in excellent agreement with the QEW universality class (see Table 1 and Fig. S7). This is consistent with the finding of Moglia et al. [40], who used a slightly more complex simulation algorithm to model the growth of cancer cell monolayers. Without obstacles, our simulations reproduce earlier findings [13] (Table 1). In particular, we find *z* = 3/2 without obstacles and *z* ≈ 1.15 at the critical obstacles density, which allows us to compute the scaling exponent of the sector boundaries from eq. (5) as *ζ* = 1*/z*. This gives *ζ* = 2/3 and *ζ* ≈ 0.86, in excellent agreement with the lineage fluctuation exponents *ζ* = 0.66 and *ζ* ≈ 0.86 in Fig. 5d. By contrast, the clone size distribution *P* (*A > a*) for bubbles is predicted to scale as *a*^-1^/(^1+*z*)^, which gives *a*^-2^/^5^ in the KPZ case and predicts *a*^-0^.^46^ for QEW. However, we observe the same scaling in our simulations both without obstacles and at the critical obstacle density (see Fig. S6b, inset).

## Supplementary results

**Figure S1.**
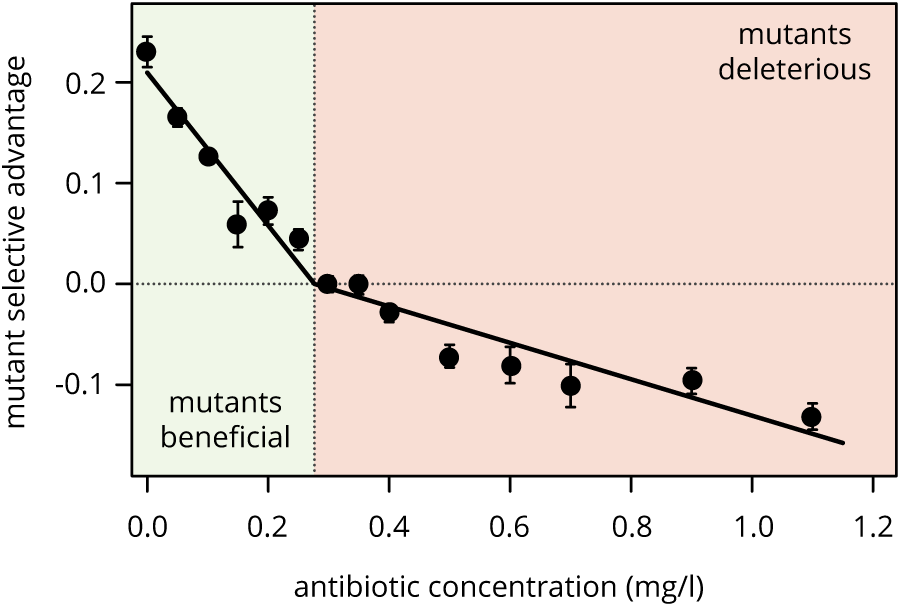
Selective difference between mutants and wild type at different concentrations of doxycycline and tetracycline. For low antibiotic concentration, plasmid loss is beneficial such that mutants have an advantage, while the mutants become first neutral and then deleterious for higher concentrations.

**Figure S2.**
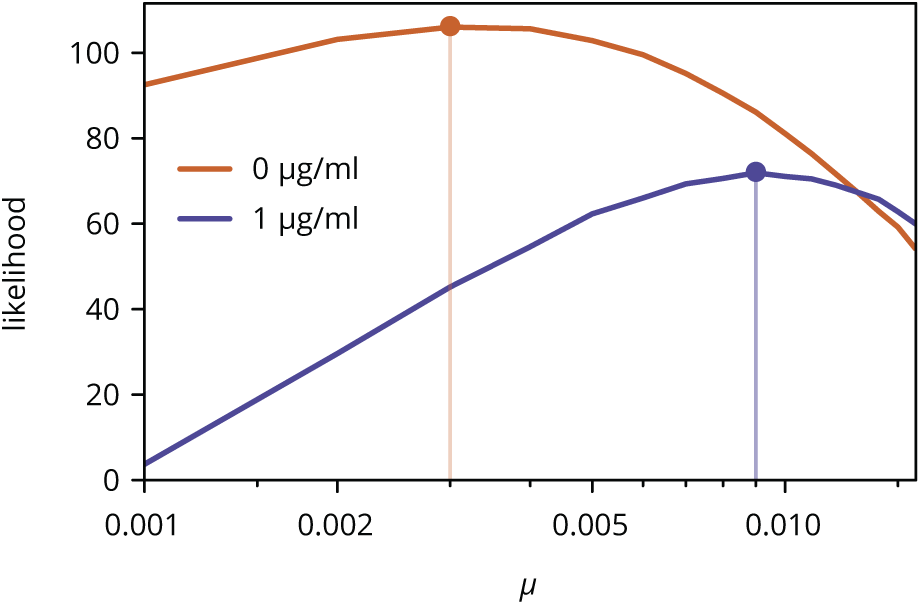
Likelihood of mutation rate for two different concentrations of doxycycline. Only the curves with the maximum likelihood value for *s* are shown.

**Figure S3.**
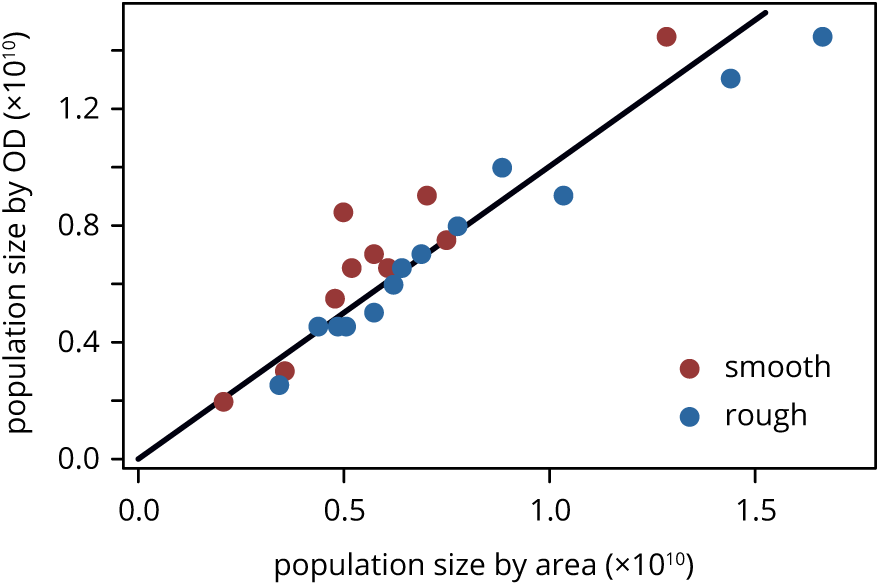
Population size of colonies estimated by measuring colony height and area, and by resuspending and measuring the resulting optical density. The black line has slope 1.

**Figure S4.**
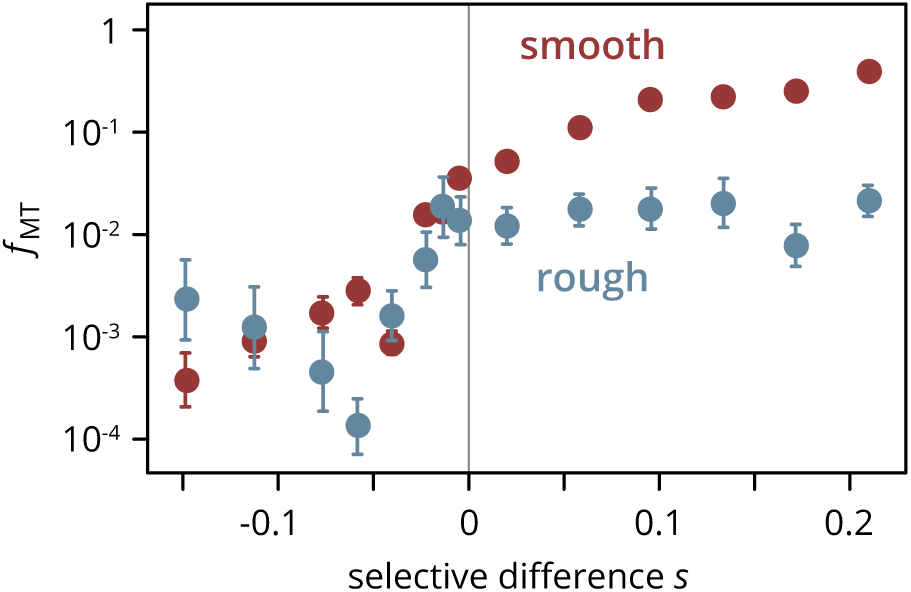
Final frequency of mutants after 3 days of growth, on smooth (red) and rough (blue) plates. Same data as Fig. 2a, plotted on a semi-logarithmic scale.

**Figure S5.**
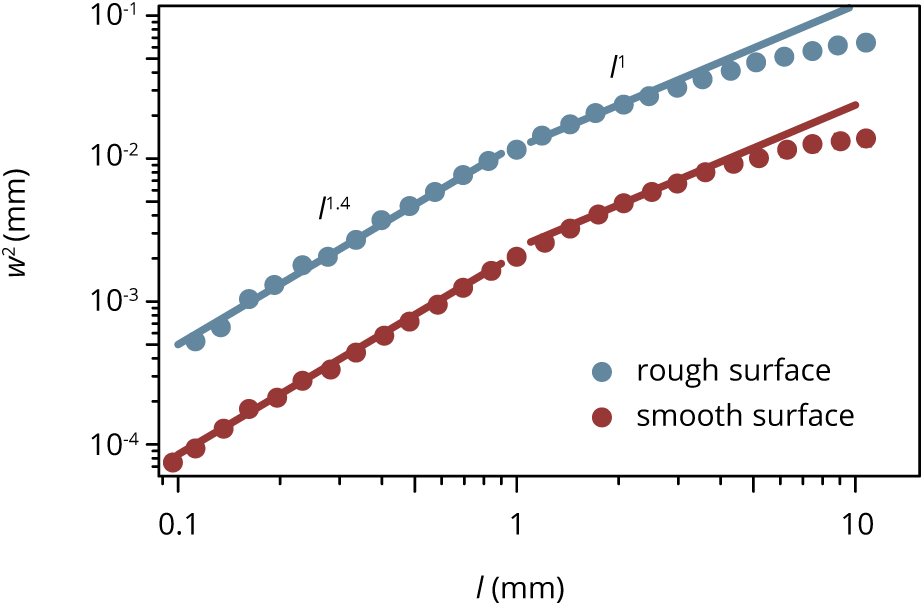
Roughness of colonies of different media, defined as the mean square deviation from the best-fit circle. Colony boundaries were extracted and the variance *w*^2^ of the radius fluctuations measured over different window lengths *l*. For interfaces in the KPZ universality class, *w*^2^ ∼ *l*.

**Figure S6.**
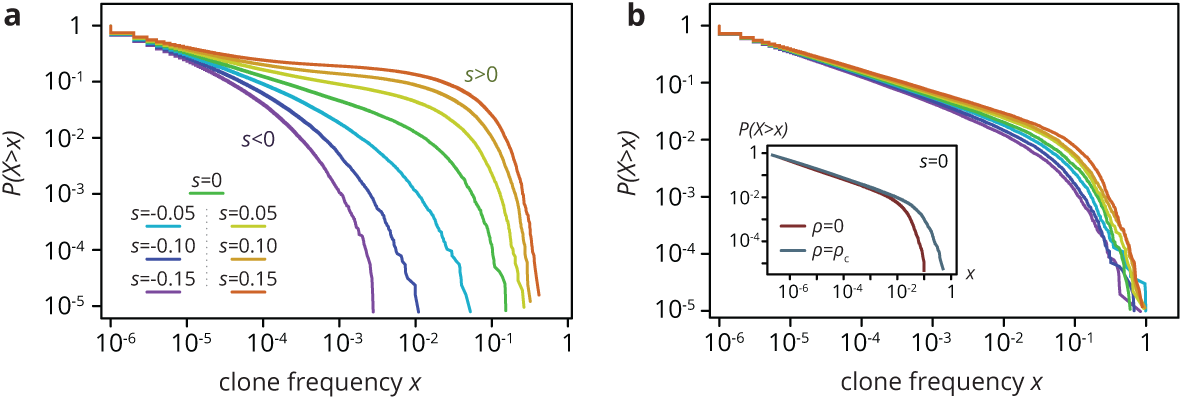
Clone size distribution *P* (*X > x*) for colonies grown on smooth (a) and rough (b) substrates in simulations. Deleterious mutants are shown in magenta tones; their clones are typically small. Neutral clones are shown green; their size distribution is broad. Advantageous mutations (red tones) are even more broadly distributed, as large sectors establish more often. In the absence of environmental disorder, the clone size distribution *P* (*X > x*) was very broad, with a low-frequency power-law regime *x*^-2/5^ corresponding to mutant bubbles and a steeper power-law at high frequencies characterizing sector sizes [16]. A deleterious fitness effect *s* of the mutations created an effective cut-off because sectors no longer formed, whereas positive *s* increased the likelihood of high-frequency clones, because sectors establish more often and grow to larger frequencies when they do. At the critical obstacle density, we found that a neutral clone size distribution that was remarkably similar to what we found without environmental heterogeneity (b, inset). By contrast, selective differences between mutants and wild type had a much less pronounced effect on the clone size distribution as the density of obstacles increased. The clone size distribution for both beneficial and deleterious mutations thus resembled the distribution for neutral mutations. In simulations at the critical obstacle density *ρ* = *ρ*_*c*_, *P* (*X > x*) became roughly independent of *s*, such that fitness effects associated with the mutations were effectively inconsequential at the level of individual clones.

**Figure S7.**
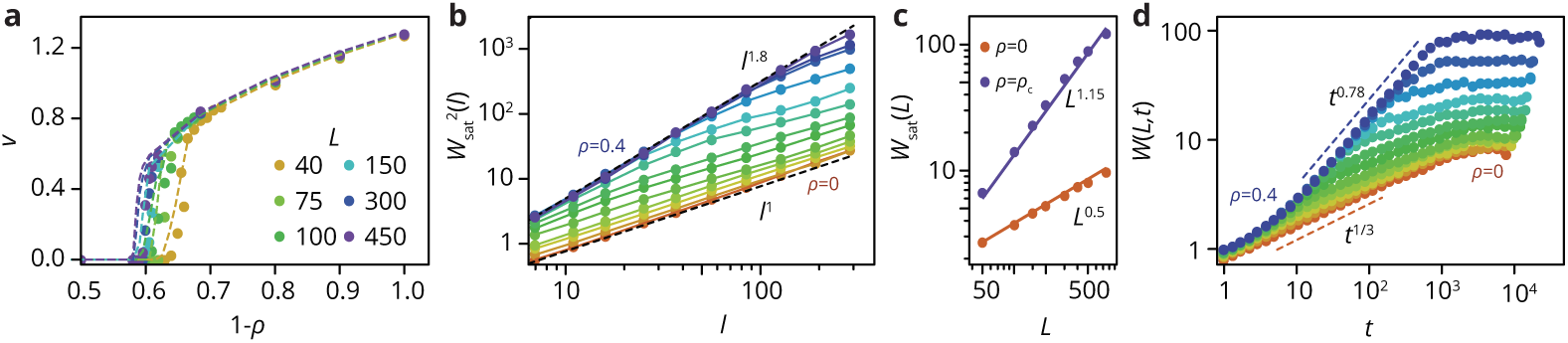
Characterization of Eden clusters with obstacles. (a) Taking the front speed *v* as the order parameter, there is a phase transition at a critical obstacle density *ρ*_*c*_(*L*) depending on system size *L*. For an infinite system, *ρ*_*c*_ ≈ 0.41 ± 0.01. For *ρ < ρ*_*c*_, we have *v* ∼ *|ρ - ρ*_*c*_*|*^0.21^. (b) Roughness *W* of fully developed interfaces at *L* = 400 for varying obstacle densities, as a function of the window length *l*, such that *W* ∼ *t*^*α*^. At *ρ* = 0, we find *α* = 1/2, consistent with the KPZ universality class. At *ρ*_*c*_, we find *α*_loc_ ≈ 0.9, consistent with the qEW universality class, which is known to be characterized by two roughness exponents, a local exponent (*α*_loc_), and a global exponent *α*_*G*_ ≈ 1.15 *> α*_loc_ when the roughness is measured over the whole system size (see panel c). The time evolution *W* (*t*) ∼ *t*^*β*^ of the interface also follows dynamics consistent with KPZ (*β* = 1/3) and qEW (*β* ≈ 0.78) in the limiting cases. For intermediate 0 ≪ *ρ ≪ ρ*_*c*_, there is a crossover from qEW at short times to KPZ dynamics at longer times, before the roughness saturates at a *ρ*-dependent value. For easier analysis, all simulations were performed in a box-like geometry.

